# Differential efficacy of α5IA in the Dp(16)1Yey mouse model of Down syndrome: implications for translational research

**DOI:** 10.64898/2026.05.12.724517

**Authors:** Jérémy Jehl, Valérie Nalesso, Claire Chevalier, Véronique Brault, Marie-Claude Potier, Elodie Ey, Yann Hérault

## Abstract

Cognitive impairments significantly impact the daily life of people with Down syndrome (DS). Overinhibition mediated by interneurons in the central nervous system was proposed as a key pathophysiological mechanism. Previous studies demonstrated cognitive rescue in the Ts65Dn mouse model using α5IA, a negative allosteric modulator of the α5 subunit-containing GABA_A_ receptors.

Here, we evaluated the effect of this drug in a mouse model carrying a more accurate duplication of the orthologous region to the human chromosome 21, namely the Dp(16)1Yey mouse model. First, we expanded the phenotypic characterization of Dp(16)1Yey mice using translationally more relevant behavioral tests. We confirmed spatial memory deficits in Dp(16)1Yey mice in the Barnes maze, and highlighted robust learning deficits in the pattern dissociation task and impairments in motor coordination. Next, we evaluated the effect of α5IA treatment on cognitive and motor performance. While α5IA treatment improved motor coordination in the Dp(16)1Yey mice, it failed to restore cognitive performance in the Barnes maze or in the pattern dissociation task.

These findings could suggest divergent pathophysiological mechanisms between the Dp(16)1Yey and the Ts65Dn models. Potentially, it could explain the limited efficacy of similar pharmacological intervention in clinical trials for DS. Further preclinical studies should prioritize refined behavioral paradigms and probably the use of more complex DS models to enhance the translational potential of candidate therapies.

## Introduction

Down syndrome (DS) is the most prevalent aneuploidy, occurring in approximately 1 in 800 births (Bull, 2020). It represents the most common genetic cause of intellectual disability (ID). While clinical manifestations vary among individuals with DS, all of them exhibit ID, typically associated with impairments in prefrontal and hippocampal functions (Grieco et al., 2015). These deficits contribute to challenges in long-term memory, working memory, and general learning abilities, often leading to difficulties in social integration (Antonarakis et al., 2020; Grieco et al., 2015). ID in this syndrome originated during neurodevelopment, with structural brain differences observed in both humans and mouse models compared to the normotypic population (Antonarakis, 2017; Herault et al., 2017). Beyond cognitive impairments, DS is also characterized by a range of clinical features, including hypotonia, delayed motor development, gait and balance abnormalities, congenital heart defects and craniofacial malformations (Antonarakis et al., 2020; Bull, 2020), reflecting the multisystemic nature of the syndrome.

Modeling DS is essential for understanding its underlying mechanisms and for evaluating potential therapeutic strategies. This syndrome presents a unique challenge due to the complexity of its genetic basis, involving the triplication of 233 protein-coding genes, 423 non-coding genes, and 188 pseudogenes. In mice, regions homologous to human chromosome 21 (Hsa21) are spread across three chromosomes: Mmu10, Mmu16, and Mmu17, with the largest syntenic region located on Mmu16 (Herault et al., 2017).

Since its development (Davisson et al., 1990, 1993; Reeves et al., 1995), the Ts65Dn model has been the gold standard for studying DS-related phenotypes and evaluating preclinical therapies (Herault et al., 2017). However, this model has notable limitations. While it carries an extra minichromosome with a substantial portion of the Mmu16 region syntenic to Hsa21, non-syntenic expressed genes are also triplicated (Duchon et al., 2011, 2022).

The Dp(16)1Yey mouse model was developed in 2007, featuring a complete segmental duplication of the Mmu16 syntenic region without additional genetic material (Li et al., 2007). The Dp(16)1Yey model exhibits key DS hallmarks, including cardiac malformations, craniofacial abnormalities, and cognitive deficits (Duchon et al., 2021; Li et al., 2007). Notably, both Ts65Dn and Dp(16)1Yey mice display memory impairments reminiscent of those observed in individuals with DS (Duchon et al., 2021; Goodliffe et al., 2016).

One mechanism involved in DS-related cognitive deficits is the disruption of the excitatory/inhibitory (E/I) balance in the brain (Fernandez et al., 2007; Kleschevnikov et al., 2004; Rueda et al., 2008), a common feature in various neurological disorders (Marín, 2012; Ramamoorthi & Lin, 2011). Further studies using DS mouse models have revealed an E/I imbalance skewed toward inhibition, as demonstrated by molecular and electrophysiological analyses (Kurt et al., 2000; Zorrilla de San Martin et al., 2018, 2020).

Restoring the E/I balance was a promising therapeutic strategy for DS. One approach involves modulating the activity of GABA receptors, particularly the GABA_A_ subtype. Early studies using low doses of pentylenetetrazol (PTZ), a non-selective GABA_A_ receptor antagonist, demonstrated that treatment in Ts65Dn mice leads to improvements in declarative and spatial memory deficits (Fernandez et al., 2007; Rueda et al., 2008). Therefore, despite its withdrawal from clinical use due to its pro-convulsant properties, PTZ helped validate the concept of targeting GABAergic inhibition to alleviate cognitive deficits in DS.

To circumvent these adverse effects, the α5 subunit-containing GABA_A_ receptors (GABA_A_ α5) was a promising candidate. These receptors account for only ∼5% of total GABA_A_ receptors and are predominantly expressed in the hippocampus and prefrontal cortex, two regions central to DS-related cognitive impairments (Mohamad & Has, 2019; Sieghart & Sperk, 2002). Notably, mice lacking GABA_A_ α5 subunits exhibit enhanced spatial memory (Collinson et al., 2002), indicating that selective inhibition of these receptors improve cognition without inducing seizures.

The use of negative allosteric modulators (NAM) specific to GABA_A_ α5 receptors, such as the 3-(5-methylisoxazol-3-yl)-6-[(1-methyl-1,2, 3-triazol-4-yl)methyloxy]-1, 2, 4-triazolo[3, 4-a]phthalazine (α5IA) has shown promising results to improve cognition (Sternfeld et al., 2004). Acute α5IA treatment of Ts65Dn mice improved working, declarative, and spatial memories (Braudeau et al., 2011; Duchon et al., 2020). *In vivo* electrophysiological studies in CA1 further revealed restored long-term potentiation (LTP), a critical mechanism for learning and memory, following α5IA administration (Duchon et al., 2020). Similar findings were reported with other GABA alpha 5 NAMs (Martínez-Cué et al., 2013b).

Building on these findings, a Phase II clinical trial monitored the efficacy and safety of basmisanil (RG1662), another selective GABA_A_ α5 NAM, in adolescents and young adults with DS (Goeldner et al., 2022). While basmisanil demonstrated target engagement and a good safety profile, it failed to achieve the primary endpoint of significantly improving cognition and adaptive functioning compared to placebo that itself had a beneficial effect. These outcomes underscore the challenges of translating preclinical efficacy into clinical benefit and highlight the need for refined endpoints and potentially earlier intervention approaches.

In this study, we investigated the effects of α5IA in the Dp(16)1Yey mouse model, which carries a full duplication of the Mmu16 syntenic region to Hsa21 and more closely mirrors the DS genetic complexity. We first confirmed the presence of cognitive deficits, such as working memory and object recognition, and motor deficits in the rotarod test, as previously reported (Aziz et al., 2018; Duchon et al., 2021; Nguyen et al., 2018; Siegel et al., 2023). To enhance translational relevance, we also employed additional paradigms with the visual discrimination test using the touchscreen technology (Aziz et al., 2018; Duchon et al., 2021; Nguyen et al., 2018; Siegel et al., 2023), and spatial learning with the Barnes maze, to minimize stress-related confounders observed in the Morris water maze (Stasko & Costa, 2004). Next, we assessed the efficacy of α5IA in rescuing the observed phenotypes

## Materials and Methods

### Mouse line

The B6.129S7-Dp(16Lipi-Zbtb21)1Yey/J mice, noted here Dp(16)1Yey mice (Li et al., 2007), were obtained originally from Dr Eugene Yu more than 15 years ago in the laboratory of Dr Yann Hérault. Mice were bred in the Institut Clinique de la Souris (ICS), a facility approved for breeding and using animals for scientific purposes under the number D 67-218-40 (France). They were kept under controlled conditions of lighting (12h day/night cycles, night period: 7:00 pm-7:00 am), ambient temperature (22°C ± 2), and humidity (60 %) with nesting material and bedding, as well as food and water ad libitum. We used both sexes in our study to evaluate any sex differences regarding the phenotyping or the effect of the treatment. Mice were identified and genotyped following the procedure described in Supplementary methods.

### Ethical Assessment

The experimental approach was authorized by the Ministry of Research with the Ethical committee n°017 under the references APAFIS #41075-2023020911351260v4 and APAFIS #46837-2023111415162019v3.

### Behavioral phenotyping pipelines

Cohorts of mutant mice with their WT littermates were kept in constant groups of four (2 WT and 2 Dp(16)1Yey same-sex mice) from weaning on and throughout the test starting at the age of 8 weeks. We split the behavioral tests into two pipelines to limit “side effects” like age and to assess cognitive functions around the same age (+/− 1 week; Figure 1A). In the first pipeline (P1), we performed the Barnes maze, the rotarod and the novel object recognition (NOR). The second pipeline (P2) was composed of the pattern dissociation task (Jehl et al., 2025) and the Y-maze. Considering these two pipelines and the expected effect size, we performed power test calculation to determine the number of individuals sufficient to detect a change with 80% confidence, with a significant level of 5% (alpha error) in our treatment pipeline (Table S1).

**Figure 1:**
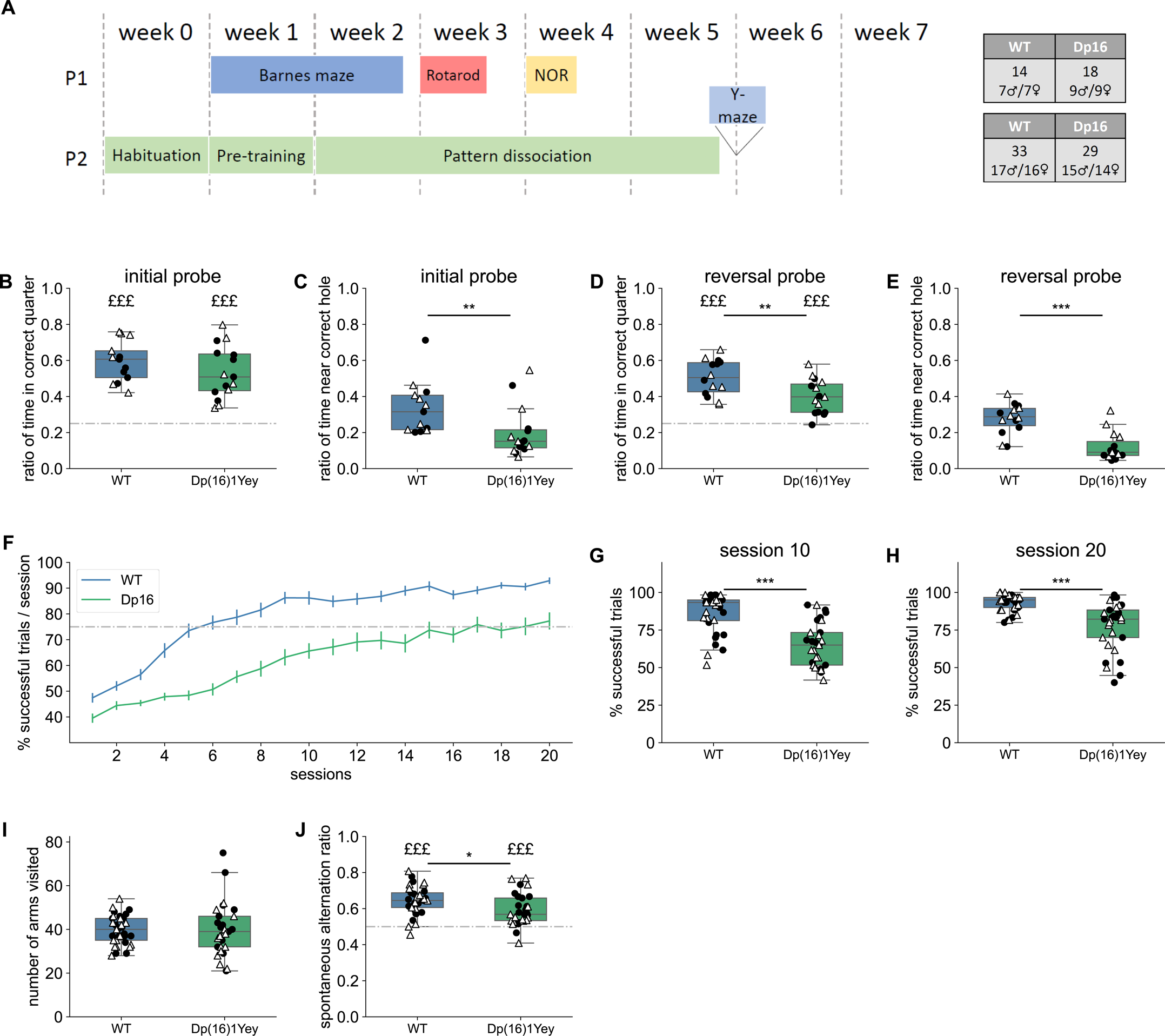
Impairments in working and spatial memory and delayed learning in Dp(16)1Yey mice. (A) Pipelines P1 and P2 were used to complement and validate the phenotypic characterization of the Dp(16)1Yey mouse model. All mice were aged of 8 weeks on week 1. The grey tables present the number of mice used in the two characterization pipelines. Ratio of time spent in the correct quarter (B) and near the correct hole (C) over the total time of exploration (90 s) during the initial probe test of the Barnes maze (n WT = 13, Dp(16)1Yey = 15). Ratio of time spent in the correct quarter (D) and near the correct hole (E) over the total time of exploration (90 s) during the reversal probe test of the Barnes maze (n WT = 14, Dp(16)1Yey = 15). The dotted lines represent random exploration (0.25) in the quarter of the Barnes maze (B, D). £££: p < 0.001 in t-tests against random exploration (B, D). *: p <0.05, **: p<0.01, ***: p<0.001 in t-tests (B, D) or u-test (C, E) between genotypes. (F). Learning curve displaying the proportion of successful trials per daily session over 20 days of learning. The horizontal dotted line represents the criterion of 75% of successful trials per session, for which we supposed mice had understood the task. Points and error bars represent mean+/-sem. Average success in the 10^th^ session (G) and the 20^th^ session (H) (n WT = 33, Dp(16)1Yey = 29). *** = p<0.001 in u-test between genotypes. (i) Number of arms visited during 8 min of free exploration of the Y-maze (t-test; n WT = 33, Dp(16)1Yey = 29). (J) Ratio of spontaneous alternations over the total alternations possible during 8 min of free exploration in the Y-maze. The dotted horizontal line represents random exploration (0.5). £££: p < 0.001 in t-tests against random exploration. * = p<0.05 in t-test between genotypes (n WT = 33, Dp(16)1Yey = 29).

Afterward, we performed a pipeline (P3) to assess the α5IA efficiency by retaining tests showing deficits in the Dp(16)1Yey mice (Figure 3A). This pipeline included the following tests: Y-maze, pattern discrimination task, Barnes maze, and rotarod. We adjusted the protocol of the Barnes maze test compared to the two previous pipelines by reducing the number of trials to attenuate any pro-cognitive effect. The full description of the experimental procedures can be found in Supplementary methods.

### α5IA treatment

For the treatment by α5IA, we used the same concentration 5mg/kg as described previously (Braudeau et al., 2011; Duchon et al., 2020). The α5IA (LS-193,268) was prepared in DMSO/Cremophor El/water (10:15:75). We performed two intra-peritoneal injections per week to each mouse starting 30 min before the first test and for the complete duration of the pipeline (6 weeks). Mice were split randomly between the treatment and the control groups. Mice received an intra-peritoneal injection of 5mg/kg of α5IA (∼25µL; treated group) or vehicle solution (∼25µL; DMSO/Cremophor El/water; control group). To balance the cage effect, each cage contained one WT treated mouse, one WT vehicle mouse, one Dp(16)1Yey treated mouse, and one Dp(16)1Yey vehicle mouse.

### Statistics

All tests were performed with 5% chance of rejecting the null hypothesis when it is actually true. For all the tests, we determined the normality of the data using the Shapiro-Wilk test. Then, variance homogeneity was tested using the Levene test as suggested by Hosken and colleagues for analyzing behavioral data (Hosken et al., 2018). For comparisons between genotypes in the first and second pipelines (Y-maze, NOR, Barnes maze, touchscreen, rotarod acceleration paradigm), data that fulfilled the normal distribution and homoscedasticity conditions were compared using the two tailed Student t-test. The rest of the data (rotarod at fixed speed) were analyzed using the Mann-Whitney U-test. When compared to specific values, data with normal distribution were evaluated using one tailed one-sample t-test or Wilcoxon signed-rank centered on our reference value if the data did not follow normal distribution.

The power test calculations for determining the number of animals for the treatment cohorts were performed using the G*power software (v3.1.9.7) that has been developed for this kind of analysis (Faul et al., 2007, 2009) (Table S1). When comparing the different groups from the treatment cohort, we performed ANOVA followed by a Tuckey post-hoc test if the data followed a normal distribution and were homogeneous; otherwise, we used Kruskal-Wallis followed by Dunn tests. We also evaluated the impact of genotype and sex using a two-way ANOVA in the phenotyping pipelines (P1 and P2). We added the factor treatment in a three-way ANOVA for the treatment pipeline (P3). We did not observe any significant sex effects unless otherwise specified. Results of statistical analyses are summarized in Tables S2 and S3.

For the multivariate analysis, we used the GDAPHEN pipeline (Muñiz Moreno et al., 2023) to identify the key variables that differentiate Dp(16)1Yey and their wild-type littermates with or without treatment with the classical selection of non-correlated variables (Pearson correlation below 0.75). To identify the most discriminant variables, we initiated the analysis with 14 variables derived from all the tests (Table S4).

## Results

### Pipeline-dependent memory deficits in Dp(16)1Yey mice: insights into DS cognitive variability

To identify robust phenotypes for preclinical studies, we reevaluated spatial memory, learning capacity and motor coordination in the Dp(16)1Yey mice with our local conditions using two pipelines (Figure 1A).

In pipeline 1 (P1; Figure 1A), a single cohort of mice, encompassing both genotypes and sexes, was first evaluated in the Barnes maze to asses spatial memory by removing the escape box after the learning period. Both WT and Dp(16)1Yey mice successfully remembered the quarter containing the escape box, with no significant difference between groups (WT vs random : t(13)=11.025, p<0.001; Dp(16)1Yey vs random : t(14)= 7.609, p<0.001; WT vs Dp(16)1Yey: t(27)=1.213, p=0.236; Figure 1B, Table S2). However, Dp(16)1Yey mice spent significantly less time near the expected escape hole compared to WT (WT vs Dp(16)1Yey: U=2.919, p=0.005), showing reduced spatial memory precision (Figure 1C).

During the reversal probe, Dp(16)1Yey mice still preferred the correct quarter (WT vs random: t(13)=9.589, p<0.001; Dp(16)1Yey vs random: t(14)=6.072, p<0.001), but to a lesser extent than WT mice (WT vs Dp(16)1Yey: t(27)=2.917, p=0.007; Figure 1D). Moreover, Dp(16)1Yey mice spent significantly less time near the expected escape box position (WT vs Dp(16)1Yey: U=192, p<0.001; Figure 1E). These results suggest that the shorter reversal learning phase was more challenging for Dp(16)1Yey mice or that these mice exhibited impairments in cognitive flexibility. A two-way ANOVA (genotype and sex factors) for the time spent near the correct hole revealed a significant sex effect (sex effect: F(1,28)=4.455, p=0.045) but no interaction between sex and genotype (interaction sex-genotype: F(1,28)=1.024, p=0.321; Table S2). Overall, compared to WT littermates, Dp(16)1Yey mice presented reduced spatial memory in the Barnes maze, and the deficit was more pronounced in the reversal phase as in previous studies in the Morris water maze (Goodliffe et al., 2016).

To evaluate motor function, mice performed the rotarod test. On the first day, mice were tested at fixed rotation speeds. In the easy challenge (16 RPM; Figure 2A) and medium challenge (24RPM; Figure 2B), Dp(16)1Yey mice succeeded in performing the 2 min trials similarly to WT mice (WT vs Dp(16)1Yey at 16 RPM: U=120.5, p=0.780; 24 RPM: U=166.5, p=0.081). However, when Dp(16)1Yey mice were placed in the most challenging conditions (32 RPM; Figure 2C), they fell significantly earlier than their WT littermates (WT vs Dp(16)1Yey: U=208, p=0.002). These results were confirmed during the second day, when the mice performed the rotarod test with an acceleration ranging from 4 to 40 RPM over 5 min. In this case, Dp(16)1Yey mice also fell significantly earlier than WT mice (WT vs Dp(16)1Yey: t(30)=3.741, p<0.001; Figure 2D, Table S2, with an average fall speed between 24 and 30 RPM. In contrast, WT mice fell on average at a speed higher than 32 RPM. These results confirmed motor impairments in the Dp(16)1Yey as shown previously (Aziz et al., 2018).

**Figure 2:**
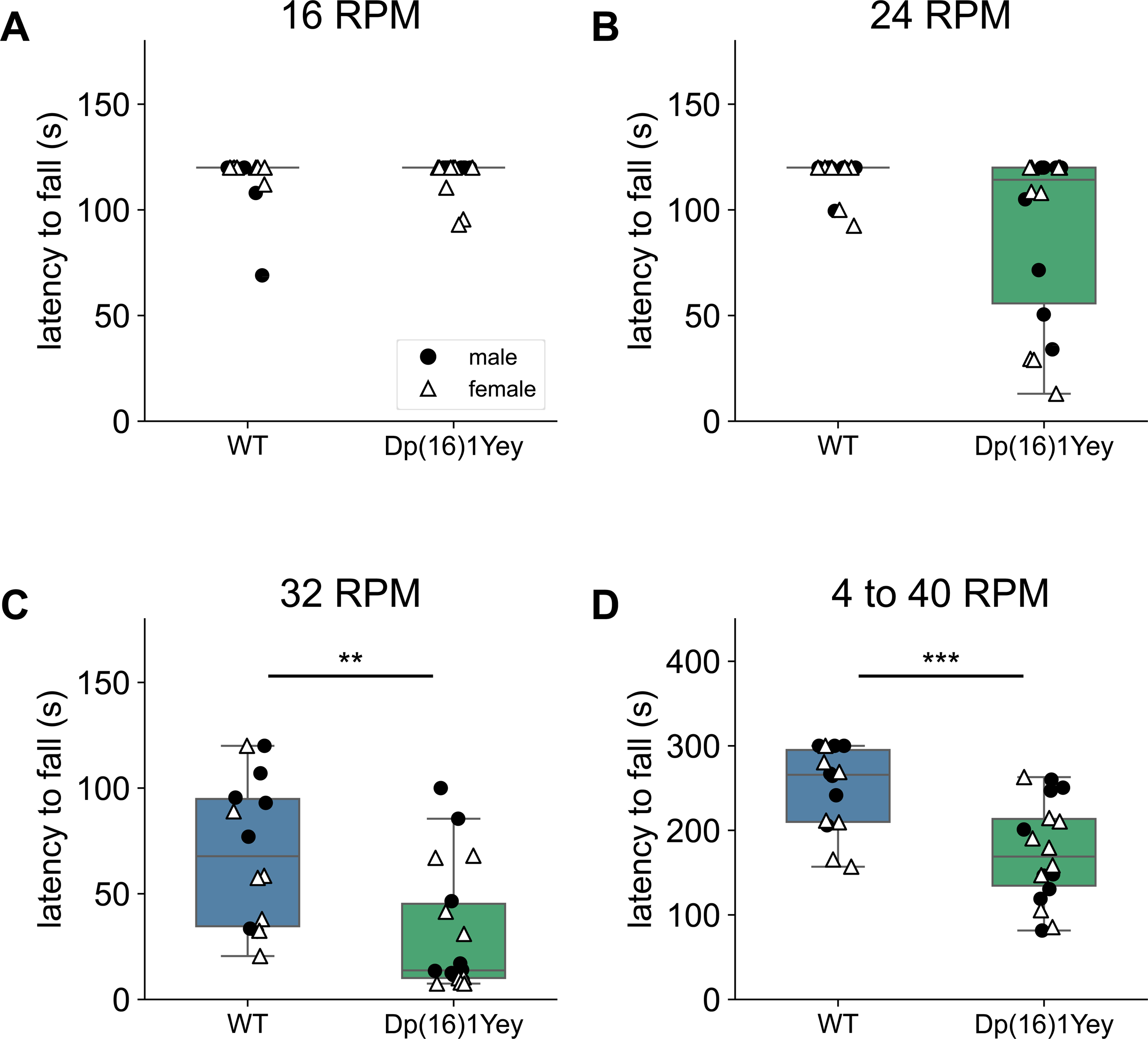
Motor coordination deficits in Dp(16)1Yey mice. Latency to fall off the rod in 2 min test with a stable rotation speed of (A) 16 rotations per minute (RPM), (B) 24 RPM, and (C) 32 RPM. (D) Latency to fall off the rod with an increasing rotation speed between 4 RPM and 40 RPM over 5 min. n WT = 14, Dp(16)1Yey = 18. **: p<0.01, ***: p<0.001 in u-tests (A-C) or t-test (D) between genotypes.

Then, we investigated the declarative memory of Dp(16)1Yey mice using a novel object recognition test with 24-hour retention interval between familiar object acquisition and novel object presentation. During the acquisition phase, both genotypes explored the objects for similar duration and display no side preference (Figure S1C,D; Table S2). After the retention period, both WT and Dp(16)1Yey mice demonstrated a significant preference for the novel object over the familiar one (both groups t-test against randomness (0.5): p<0.01) with no significant difference emerged between them in their ratio of exploration (t-test WT vs Dp(16)1Yey: p=0.728; Figure S1E, Table S2). Contrary to previous findings in our lab (Duchon et al., 2021), Dp(16)1Yey mice did not exhibit declarative memory deficits in the novel object recognition task. This discrepancy suggests potential interference from earlier tests within the experimental pipeline.

In the second pipeline P2 (Figure 1A), an independent cohort of mice, was tested in the touchscreen-based pattern dissociation paradigm, to learn discrimination of a positively reinforced image. WT mice reached the success threshold (75% success) by the 6^th^ training session and maintained a performance plateau at 90% accuracy from the end of the 14^th^ session onward (Figure 1F, Table S2). In contrast, the Dp(16)1Yey mice exhibited a slower improvement in correct responses (Figure 1F) and performed significantly worse than their WT littermates at both the 10^th^ session (WT vs Dp(16)1Yey: U=798, p<0.001; Figure 1G) and the 20^th^ session (WT vs Dp(16)1Yey: U=793.5, p<0.001; Figure 1H). These results highlight persistent learning deficits in the pattern dissociation task for Dp(16)1Yey mice.

After, working memory was assessed via spontaneous alternation in the Y maze test. Both wild-type (WT) and Dp(16)1Yey mice displayed comparable activity level indicated by the number of visited arms (WT vs Dp(16)1Yey: t(60)=-0.065, p=0.948; Figure 1I). Both genotypes also performed more alternations than expected by chance indicating non-random exploration (WT vs random: t(32)=10.401, p<0.001; Dp(16)1Yey vs random : t(28)=5.636, p<0.001; Figure 1J). However, the alternation ratio was lower in Dp(16)1Yey compared to WT, suggesting reduced spatial working memory efficiency (WT vs Dp16: t(60)=2.438, p=0.018; Figure 1J).

This evaluation across two experimental pipelines confirmed robust and distinct cognitive and motor phenotypes in Dp(16)1Yey mice. While spatial memory precision and motor coordination are impaired, declarative memory appears not affected, potentially due to task interference. The persistent learning deficits observed in the pattern dissociation task and reduced working memory efficiency in the Y maze underscore the complexity of cognitive dysfunction in this model. These findings highlight the importance of refined behavioral paradigms and pipeline design for accurate phenotypic characterization in DS preclinical studies.

### α5IA treatment does not change working memory in Dp(16)1Yey mice but increases exploratory behavior

Based on the results from previous pipelines, we selected tests that revealed robust deficits in Dp(16)1Yey mice to assess the efficiency of α5IA treatment. Therefore, the pipeline (P3) included the Y-maze, pattern dissociation task, Barnes maze and the rotarod test (Figure 3A).

**Figure 3:**
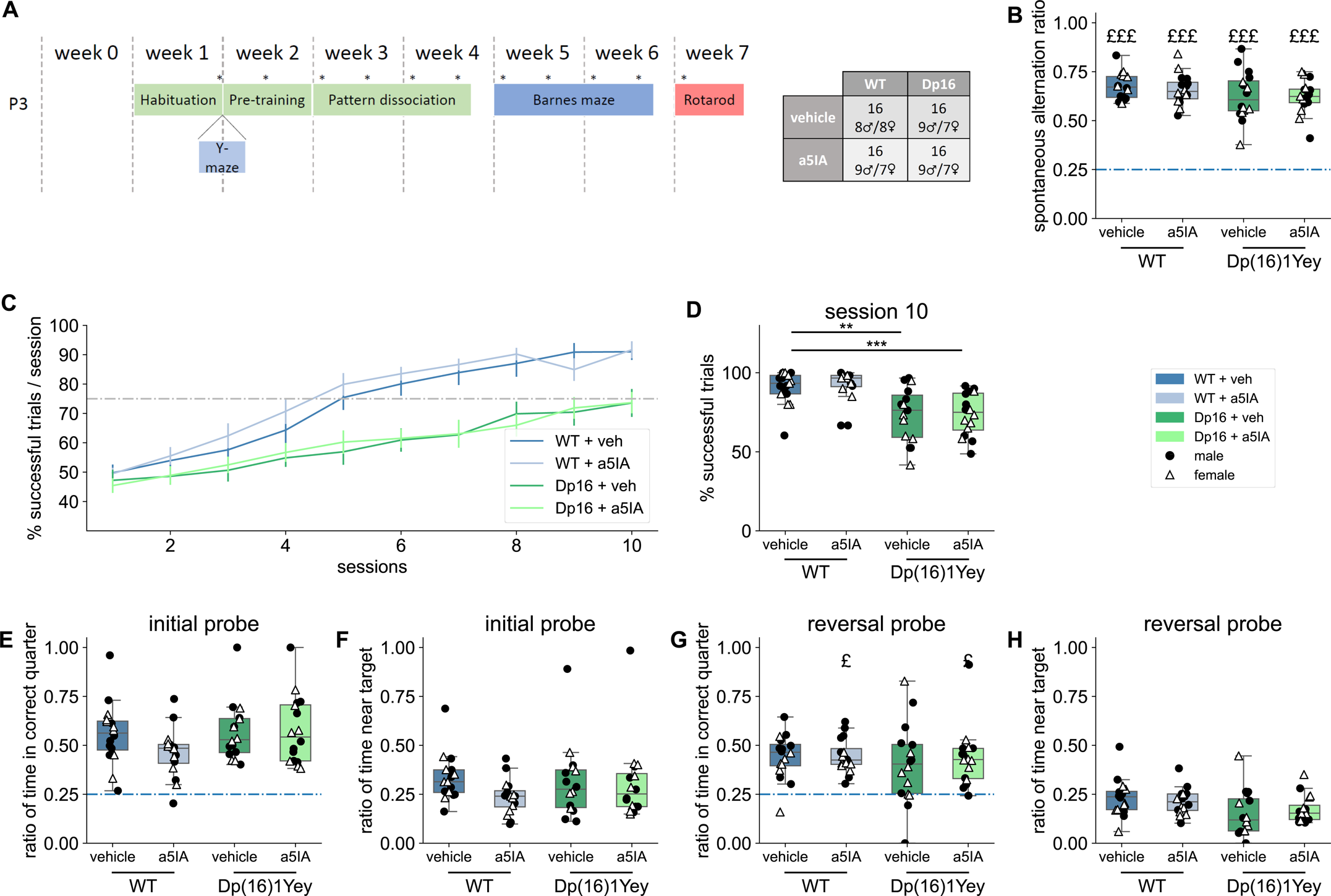
Absence of rescue of working and spatial memory deficits and delayed learning by α5IA in Dp(16)1Yey mice. (A) Pipeline P3 was use to assess the effects of the treatment in which mice received two injections per day of α5IA (* = injection of either α5IA (5mg/kg) or vehicle solution). All mice were aged of 8 weeks on Week 1. The table indicates the number of mice per group. (B) Spontaneous alternations ratio over the total alternations possible during 8 min of free exploration in the Y-maze. The dotted horizontal line represents random exploration (0.5); £££: p < 0.001 in t-test against random exploration. (C) Learning curve displaying the proportion of successful trials per daily session over 10 sessions of learning. Points and error bars represent mean+/-sem. The horizontal dotted line represents the criterion of 75% of successful trials per session, for which we supposed mice had understood the task. (D) Average success on the 10^th^ session (n = 16 in each group). **: p<0.01, ***: p<0.001 in Dunn post-hoc tests. Ratio of time spent in the correct quarter (E) and near the correct hole (F) over the total time of exploration (90 s) of the probe test of the Barnes maze. Ratio of time spent in the correct quarter (G) and near the correct hole (H) over the total time of exploration (90 s) of the reversal probe test of the Barnes maze (n=16 for each group). The horizontal dotted line represents random exploration (0.25) in the quarters. £: p<0.05 in t-test against random exploration.

In the Y-maze, both vehicle-treated groups exhibited non-random exploration (t(15)=7.179, p<0.001; t(15)=5.396, p=0.037; Figure 3B), as did the treated groups (WT treated: t(15)=5.396, p<0.001; Dp(16)1Yey treated: t(15)=3.139, p=0.003; Figure 3B). However, no significant genotype effect was detected in the percentage of spontaneous alternations, despite a trend toward reduced directed exploration in Dp(16)1Yey mice (F(3,60)=2.421, p=0.075). Unlike previous studies using α5IA (Braudeau et al., 2011; Duchon et al., 2020), no treatment effect was observed in this experiment (F(1,60)=0.031, p=0.862).

Analysis of the number of arms explored (Figure S2) revealed a sex effect (F(1,60)=5.232, p=0.026), with females exploring more arms, consistent with existing literature (Tucker et al., 2016). A significant treatment effect was also found (F(1,60)=8.668, p=0.004), though no genotype effect was observed (F(1,60)=1.311, p=0.257). Treated mice explored more arms than controls (K(3)=9.58, p=0.022), an effect that reached significance in Dp(16)1Yey mice (Post-hoc Dunn test: Dp(16)1Yey vehicle vs Dp(16)1Yey treated, p=0.029) but not in WT mice (WT vehicle vs WT treated, p=0.058; Figure S2).

In the pattern dissociation paradigm, WT vehicle-treated mice achieved the performance threshold (75% correct responses) within five training sessions (Figure 3C). However, both Dp(16)1Yey treated and Dp(16)1Yey vehicle mice failed to reach this threshold within the allocated time, consistent with earlier observations in the characterization pipeline 2 (Figure 3C). A three-way ANOVA revealed a significant genotype effect (F(1,60)=28.839, p<0.001; Table S3), but no treatment effect (F(1,60)=0.011, p=0.917) when analysing average success rates during the 10^th^ session (Figure 3D). Thus, while the learning deficit in Dp(16)1Yey mice was confirmed, α5IA treatment did not improve their performance.

We next evaluated the impact of α5IA treatment on spatial memory using the Barnes maze test. Across both probe sessions, no variable displayed a significant treatment effect (reversal probe three-way ANOVA; all treatment p-values > 0.05; Table S3). During the initial probe session, all groups preferred the correct quadrant of the maze (K(3)=4.868, p=0.181; Table S3), with each group performing significantly above chance (U-test vs. randomness, p<0.001; Figure 3E). Similarly, all groups spent a comparable proportion of time near the target hole (K(3)=5.514, p=0.138; Figure 3F). In the reversal probe session, all groups spent significantly more time than expected by chance in the target quadrant, with no difference between groups (K(3)=1.599, p=0.660; Figure 3G, Table S3). However, the time spent near the target hole varied significantly across groups (K(3)=9.061, p=0.028; Figure 3H). Post-hoc Dunn tests revealed no difference between WT vehicle and WT treated mice (p=0.582), or between Dp(16)Yey vehicle and Dp(16)Yey treated (p=0.556). Notably, both Dp(16)1Yey groups spent significantly less time near the target hole compared to WT vehicle mice (WT vehicle vs. Dp(16)1Yey vehicle: p=0.008; WT vehicle vs. Dp(16)1Yey treated: p=0.042). These results confirm the previously characterized spatial learning deficits in Dp(16)1Yey from pipeline P1 and indicate that α5IA treatment failed to rescue this impairment.

In the last rotarod test at fixed speeds, Dp(16)1Yey vehicle mice tended to show difficulty completing the slow-speed challenge (16 RPM) compared to WT vehicle mice (K(3)=6.439, p=0.092) whereas most Dp(16)1Yey treated mice successfully completed the full trial duration (Figure 4A). At medium-speed (24 RPM), Dp(16)1Yey vehicle mice performed significantly worse than WT vehicle mice (K(3)=27.748, p<0.001; Dunn test: WT vehicle vs. Dp(16)1Yey vehicle: p<0.001; Figure 4B). α5IA treatment led to a modest improvement, with Dp(16)1Yey treated mice remaining on the rod significantly longer than Dp(16)1Yey vehicle mice (Dunn test: p=0.044), though their performance still lag behind WT vehicle mice (WT vehicle vs. Dp(16)1Yey treated: p=0.026). During the high-speed challenge (32 RPM), both Dp(16)1Yey vehicle and treated mice fell significantly earlier than WT vehicle mice (K(3)=39.512, p<0.001; Dunn tests: WT vehicle vs. Dp(16)1Yey vehicle: p<0.001; WT vehicle vs. Dp(16)1Yey treated: p<0.001; Figure 4C). In the accelerating rotarod test (4 to 40 RPM over 5 minutes), a two-way ANOVA revealed significant effects of genotype (F(1,60)=54.374, p<0.001) and treatment (F(1,60)=5.171, p=0.027; Figure 4D, Table S3).

**Figure 4:**
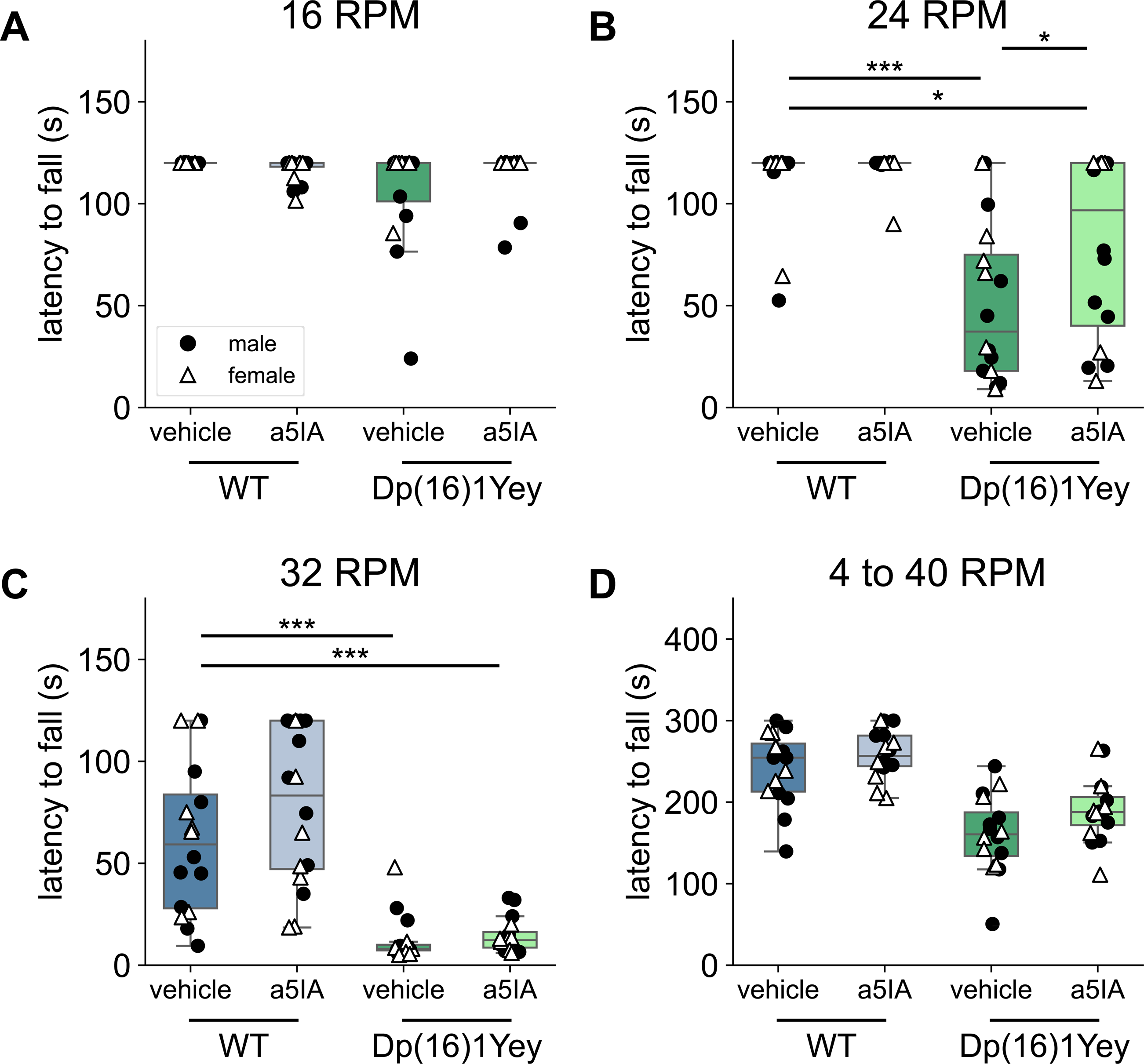
Improvement of motor impairments by α5IA in Dp(16)1Yey mice. Latency to fall off the rod in 2 min test with a stable rotation speed of (A) 16 rotations per minute (RPM), (B) 24 RPM, and (C) 32 RPM. (D) Latency to fall off the rod with an increasing rotation speed between 4 RPM and 40 RPM over 5 min. n =16 in each group. *: p<0.05, ***: p<0.001 in Dunn (B, C) or Tukey (D) post-hoc tests.

Both Dp(16)1Yey vehicle and treated mice exhibited significantly shorter latencies to fall compared to WT vehicle mice (F(3,60)=19.985, p<0.001; Tukey tests: WT vehicle vs. Dp(16)1Yey vehicle: p=0.001; WT vehicle vs. Dp(16)1Yey treated: p=0.003). However, no significant differences were observed between vehicle and treated mice within each genotype (Tukey tests: WT vehicle vs. WT treated: p=0.640; Dp(16)1Yey vehicle vs. Dp(16)1Yey treated: p=0.178). Finally, body weight measurements at the end of the test showed no significant treatment-related weight loss (F(1,60)=0.029, p=0.865; Figure S3).

### Multivariate Analysis Reveals Selective Motor Rescue by α5IA Treatment in Dp(16)1Yey Mice

To comprehensively assess the impact of genotype and α5IA treatment on behavioral phenotypes, we conducted a multivariate analysis using GDAPHEN (Genotype Discrimination using Phenotypic Features), an R-based pipeline designed to identify the most predictive qualitative and quantitative variables for genotype and treatment classification in the phenotypic datasets from pipeline P3 (Muñiz Moreno et al., 2023). This approach allowed us to integrate multiple behavioral metrics and disentangle the relative contributions of genotype, treatment, and their interactions across the dataset.

From the 14 variables initially assessed across all genotypes and treatment groups, 11 non-correlated variables were retained for discriminant analysis to avoid redundancy and ensure robustness. The first three dimensions of the discriminant space accounted for approximately 50% of the total variance (Figure 5A), providing a meaningful reduction of data complexity while preserving key phenotypic differences. The latency to fall at 24 RPM (Rpm_24), the latency to fall at 32 RPM (Rpm_32), the latency to fall during the acceleration test (acceleration_rotarod) along with the area under the learning curve (AUC_touchscreen) emerged as the primary contributors to Dimension 1, reflecting motor coordination and cognitive performance. Weight and activity in Y maze (number_arms) were the most influential variables (positive and negative, respectively) to Dimension 2 (Figure 5B), highlighting exploratory behavior and physiological differences. Contributions to Dimension 3 were more distributed, suggesting it captured residual or mixed phenotypic variation (Figure 5B).

**Figure 5:**
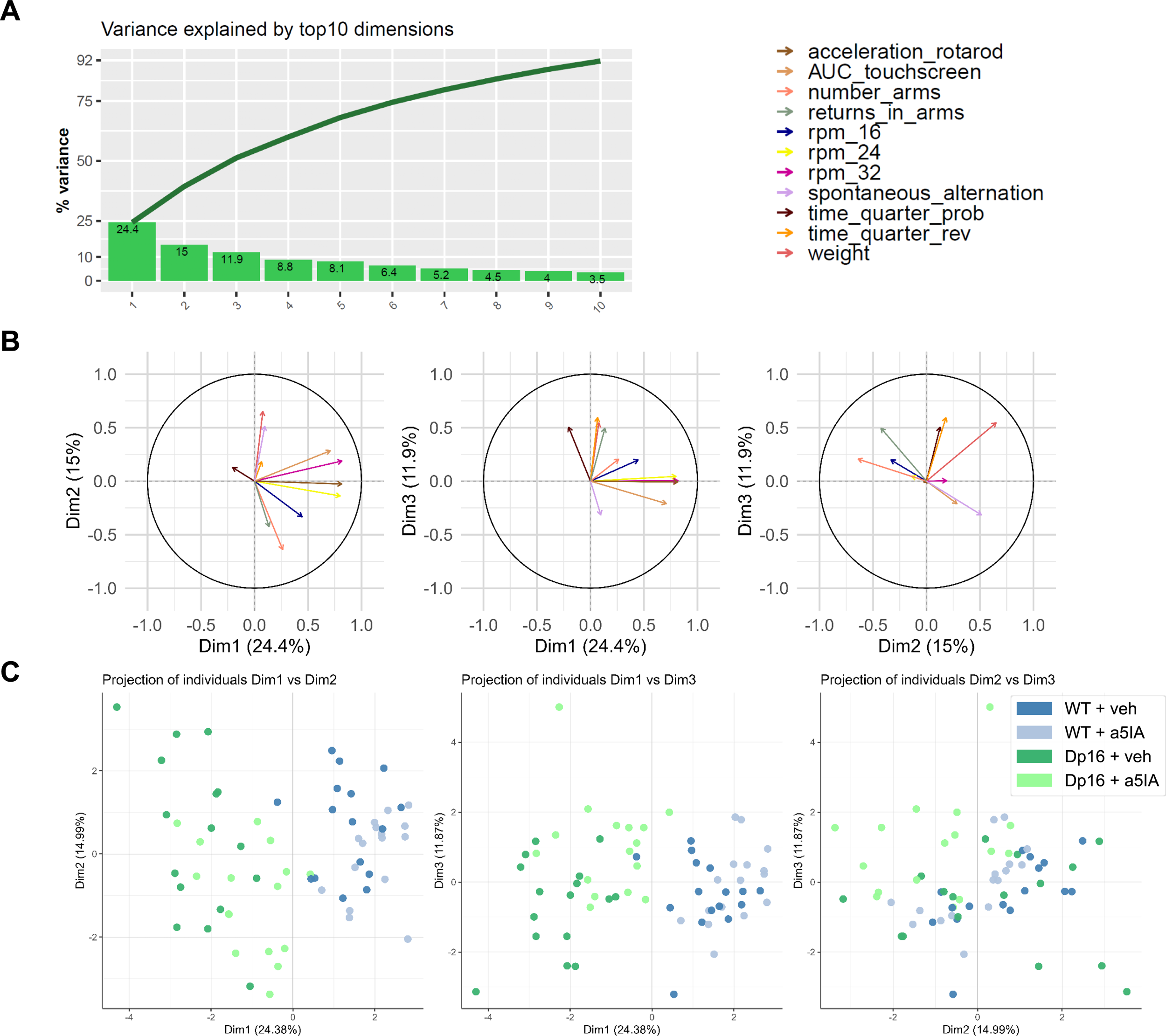
Genotype-treatment effects of α5IA in Dp(16)1Yey mouse model uncovered by Gdaphen multivariate analysis. (A) Total variance explained by the top 10 dimensions. (B) Covariance circle for the first 3 dimensions showing the contribution of each variable to each dimension. (C) 2D-PCA plots for the first 3 dimensions showing the individual animals clustering based on the PCA analyses performed with 11 non-correlated phenotypic variables derived from the 14 original behavioral variables.

Critically, the discriminant analysis revealed that Dp(16)1Yey treated mice clustered more closely with WT vehicle mice and were distinctly separated from Dp(16)1Yey vehicle mice along Dimension 1(Figure 5C). This shift indicates a partial normalization of motor-related phenotypes following α5IA treatment, aligning Dp(16)1Yey mice more closely with wild-type performance. Notably, the WT treated group also exhibited a subtle shift along Dimension 1 relative to WT vehicle mice, suggesting that α5IA may induce minor behavioral changes even in the absence of trisomy.

These findings confirm that α5IA treatment selectively improves motor coordination in Dp(16)1Yey mice, as evidenced by their closer alignment with WT mice in the discriminant space. However, cognitive performance—particularly in learning tasks—remained unimproved, underscoring the differential efficacy of α5IA across behavioral domains. This analysis not only validates the motor benefits of α5IA but also highlights the need for targeted interventions to address cognitive deficits in Down syndrome models.

## Discussion

This study confirmed working memory and motor coordination impairments in Dp(16)1Yey mice, aligning with observations in other DS models and human (Antonarakis et al., 2020; Aziz et al., 2018; Bull, 2020; Duchon et al., 2021; Goodliffe et al., 2016). Additionally, Dp(16)1Yey mice exhibited reduced spatial memory precision in the Barnes maze, while Ts65Dn mice showed delayed use of spatial strategies in a modified Barnes maze (Illouz et al., 2024). Both models also demonstrated delayed learning in touchscreen tasks, aligning with cognitive impairments observed in DS, despite protocol differences (Leach & Crawley, 2018).

Spatial memory deficits, detected in the Barnes maze, were less pronounced than in the Morris water maze, likely due to the reduced stress in the Barnes maze (Gawel et al., 2019; Pitts, 2018). Spatial memory deficits can thus be highly sensitive to protocol variations, as shown previously with the effect of water temperature altering the Ts65Dn mice performance (Stasko & Costa, 2004). This variability underscores the sensitivity of spatial memory phenotypes to protocol differences, consistent with clinical heterogeneity in DS (Grieco et al., 2015; Malegiannaki et al., 2019). Surprisingly, Dp(16)1Yey mice showed no declarative memory deficits in the novel object recognition task, contrary to prior reports (Duchon et al., 2020, 2021). We hypothesize this discrepancy may stem from pro-cognitive effects of prior testing, as this task is highly sensitive to environmental and procedural factors (Cohen & Stackman Jr., 2015). Using a single cohort of mice across multiple tests in a sequential pipeline aligns with 3R principles (Reduction, Refinement, Replacement). However, the potential confounding effects of prior training on subsequent tests must be carefully evaluated to optimize the design of behavioral testing sequences.

α5IA treatment partially improved motor coordination in Dp(16)1Yey mice, particularly at low and medium speeds, though deficits persisted at higher, more challenging speeds. This partial rescue may reflect α5IA’s modulation of somatosensory pathways, given the role of GABAα5 receptors in the somatosensory cortex (Nigro et al., 2018). However, motor deficits in DS involve multiple pathways (cerebellar, muscular, somatosensory,…), limiting the treatment’s efficacy. Notably, α5IA also slightly enhanced motor function in WT mice, suggesting a broader effect on motor systems. Unlike α5IA, the GABA-A α5 inverse agonist **RO4938581** suppressed hyperactivity in Ts65Dn mice without inducing anxiety or altering motor abilities (Martínez-Cué et al., 2013a). Additionally, the **sustained efficacy of** α**5IA** over 7 weeks of bi-weekly injections, with no desensitization, aligns with previous findings showing reduced desensitization when α5 subunits are part of the receptor composition (Caraiscos et al., 2004). This reinforces its potential for long-term therapeutic applications.

This effect was not observed across all published studies, GABA-A α5 inverse agonist. Only the GABA-A α5 inverse agonist RO4938581 treatment was shown to suppress the hyperactivity observed in Ts65Dn mice without inducing anxiety or altering their motor abilities. Furthermore, the sustained efficacy of α5IA throughout the 7-week treatment period with bi-weekly injections indicates no desensitization. This aligns with previous findings showing reduced desensitization when α5 subunits are part of the receptor composition

Unlike in Ts65Dn mice (Duchon et al., 2020), α5IA did not rescue cognitive deficits in Dp(16)1Yey mice, including spatial memory (Barnes maze) and touchscreen discrimination learning. This discrepancy may stem for spatial memory may arise from genetic differences between models and background effect (Duchon et al., 2022; Herault et al., 2017), or task-specific mechanisms. For instance, the striatum, critical for associative learning in touchscreen tasks (Delotterie et al., 2015), lacks significant GABAα5 expression, potentially limiting α5IA’s cognitive impact. This aligns with recent single-nucleus RNA-seq data (Feng et al., 2025) which reveal that DS pathology extends beyond hippocampal and cortical neurons, emphasizing the need to characterize subcortical circuits, including striatal inhibitory neuron clusters, for a comprehensive understanding of DS brain phenotypes.

Notably, other GABAα5-negative allosteric modulators (NAMs), such as RO4938581 and basmisanil, were developed to address DS cognitive impairments (Hipp et al., 2021; Martínez-Cué et al., 2013b). While RO4938581 restored spatial memory in Ts65Dn mice, the second-generation NAM basmisanil, with improved pharmacokinetics, enhanced learning and memory in the Morris Water Maze (MWM) task and rescued long-term potentiation (LTP) deficits in Ts65Dn mice(Hipp et al., 2021). However, Basmisanil ultimately failed to demonstrate cognitive benefits in clinical trials (Goeldner et al., 2022). The failure of α5IA to restore cognition in Dp(16)1Yey mice mirrors the lack of efficacy observed with basmisanil in clinical trials group (Goeldner et al., 2022). This suggests that pathophysiological differences between DS models may underlie the translational gap. Additionally, cognitive stimulation (e.g., touchscreen tasks) improved Barnes maze performance in Dp(16)1Yey mice (Figure S4), highlighting the need to account for priming effects in preclinical and clinical studies.

Unlike the robust effects observed in the Ts65dn mice, α5IA treatment in Dp(16)1Yey mice yielded only partial improvement in motor coordination, while spatial memory and learning appeared largely unaffected. These results suggest that, while GABA_A_ α5 modulation may hold promise for improving motor function in DS individuals, its cognitive benefits may depend on the genetic context and phenotypic severity across different DS models.However, several limitations must be considered: the absence of a direct comparison with the Ts65Dn model, the lack of post-treatment plasma drug level measurements, and the need for a head-to-head comparison with basmisanil/ RO4938581 to fully assess relative efficacy in a more complete DS model.

## Acknowledgements

We thank Sophie Brignon, Enzo Fontaine and Thibaut Alletto from the mouse breeding facility. We also thank Chaouki Bam’Hamed, Hamid Ennah and Mourad Korchi for animal care in the mouse behavioral facility. We are grateful to Fabrice Riet and Aline Simonet for helpful guidance for the behavioral protocols and guidance with technical issues with behavioral equipments. We thank Victorine Artot and Jean-Baptiste Oswald for their helpful comments on previous versions of the manuscript, and other members of the team for insightful discussions throughout the project.

