## Supplementary Figure for "Differential efficacy of α5IA in the Dp(16)1Yey mouse model of Down syndrome: implications for translational research"

**Supplementary figures**

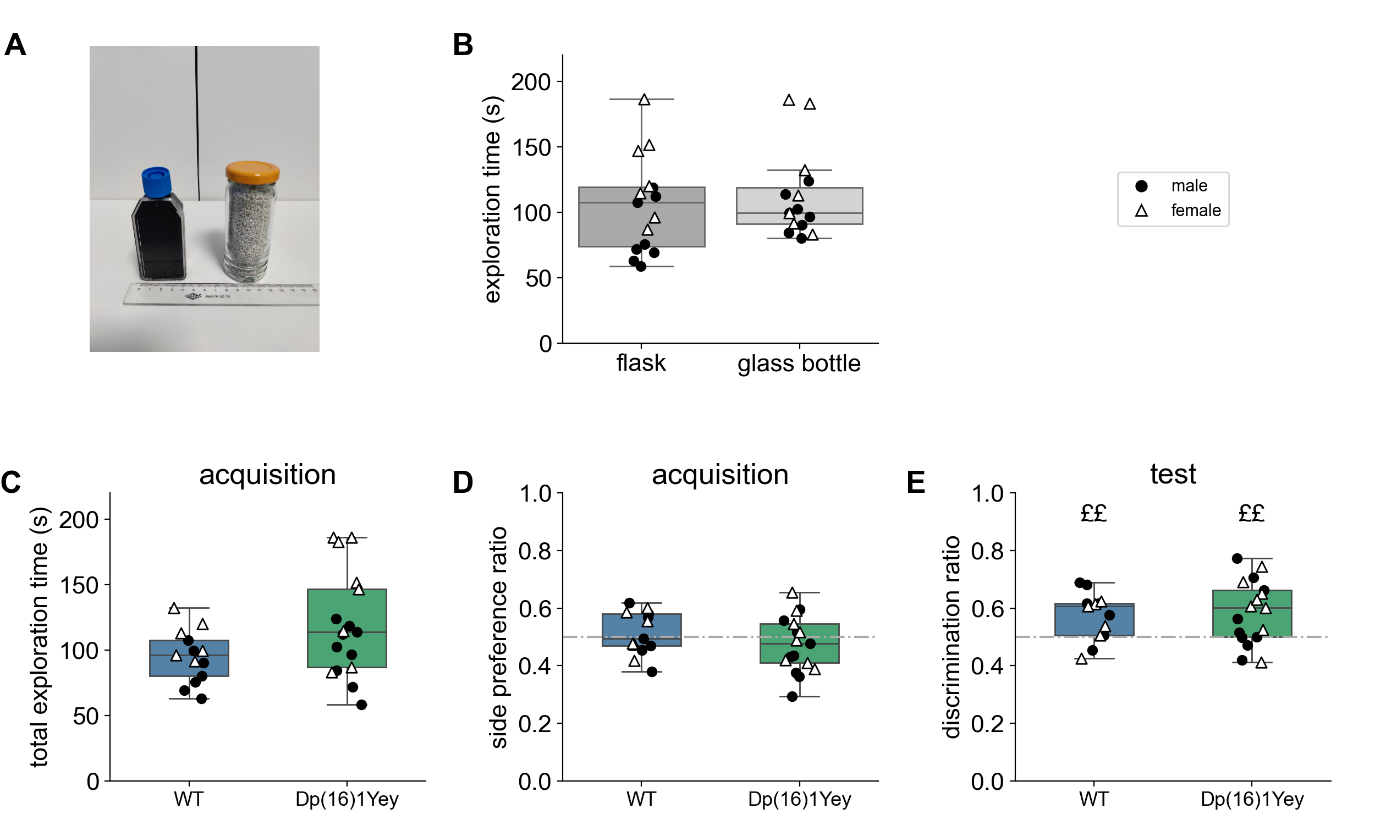

**Supplementary Figure S1: Dp(16)1Yey mice displayed typical declarative memory**. (A) The flask (left) and the glass bottle (right) that were used as objects in the novel object recognition test. (B) Similar total time of exploration between the two types of objects. Exploration was measured during the acquisition phase in which two similar objects were presented to mice. n bottle = 15, flask = 15. Genotypes and sexes were distributed evenly between the two conditions. (C) Total exploration time of the two similar objects during acquisition; no significant difference emerged between genotypes. (D) Preference index computed using the right object as a reference during the acquisition phase, in which mice were presented with two identical objects. (E) Preference index computed with the novel object as a reference during the test, in which mice were presented with two objects, one of which was different from the familiar one used in the acquisition phase. n WT = 13, Dp(16)1Yey = 17. One-tail upper t-test: ££: p<0.01.

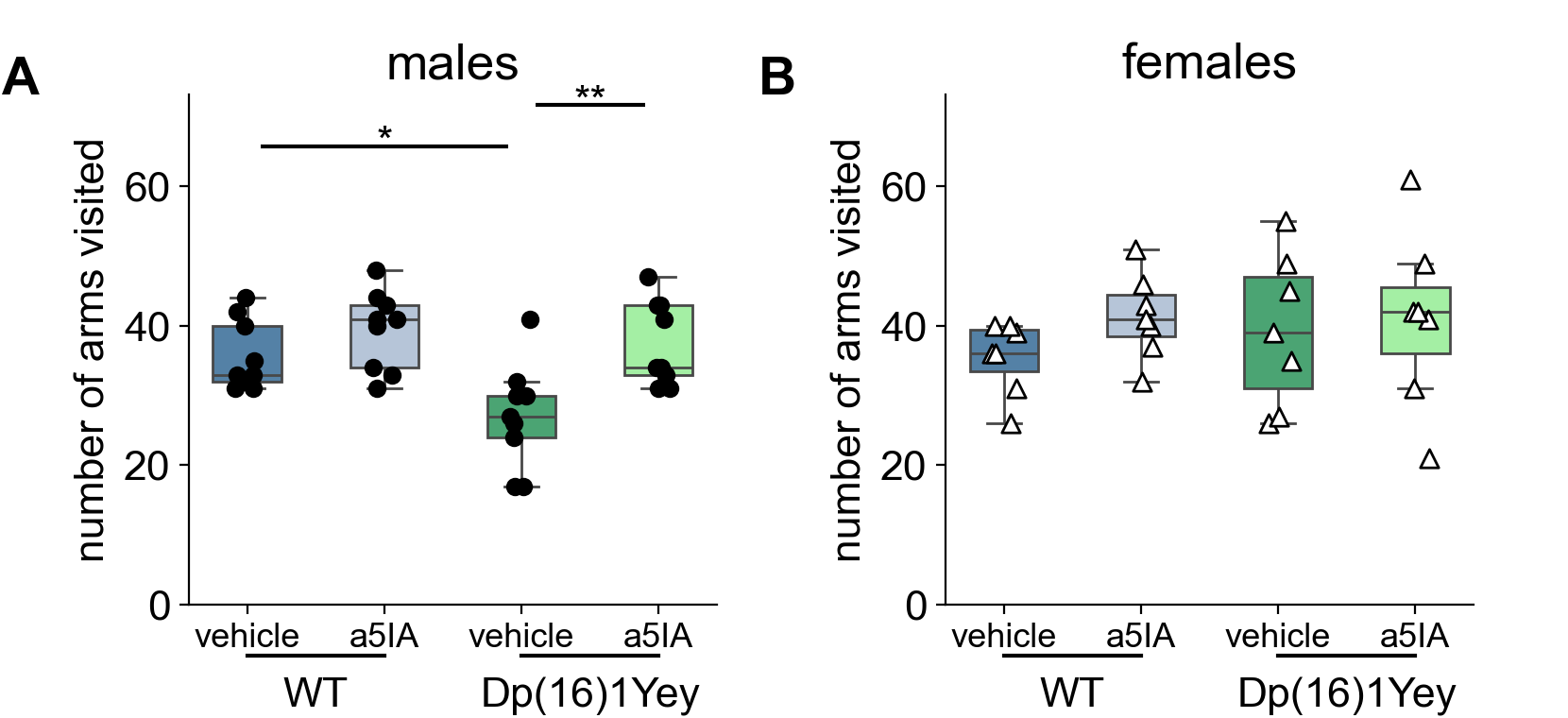

**Supplementary Figure S2: Dp(16)1Yey male mice explored less arms in the Y maze.** Number of arms visited during 8 min of free exploration of the Y-maze for (A) male and (B) female mice. * p<0.05 Tukey post-hoc test.

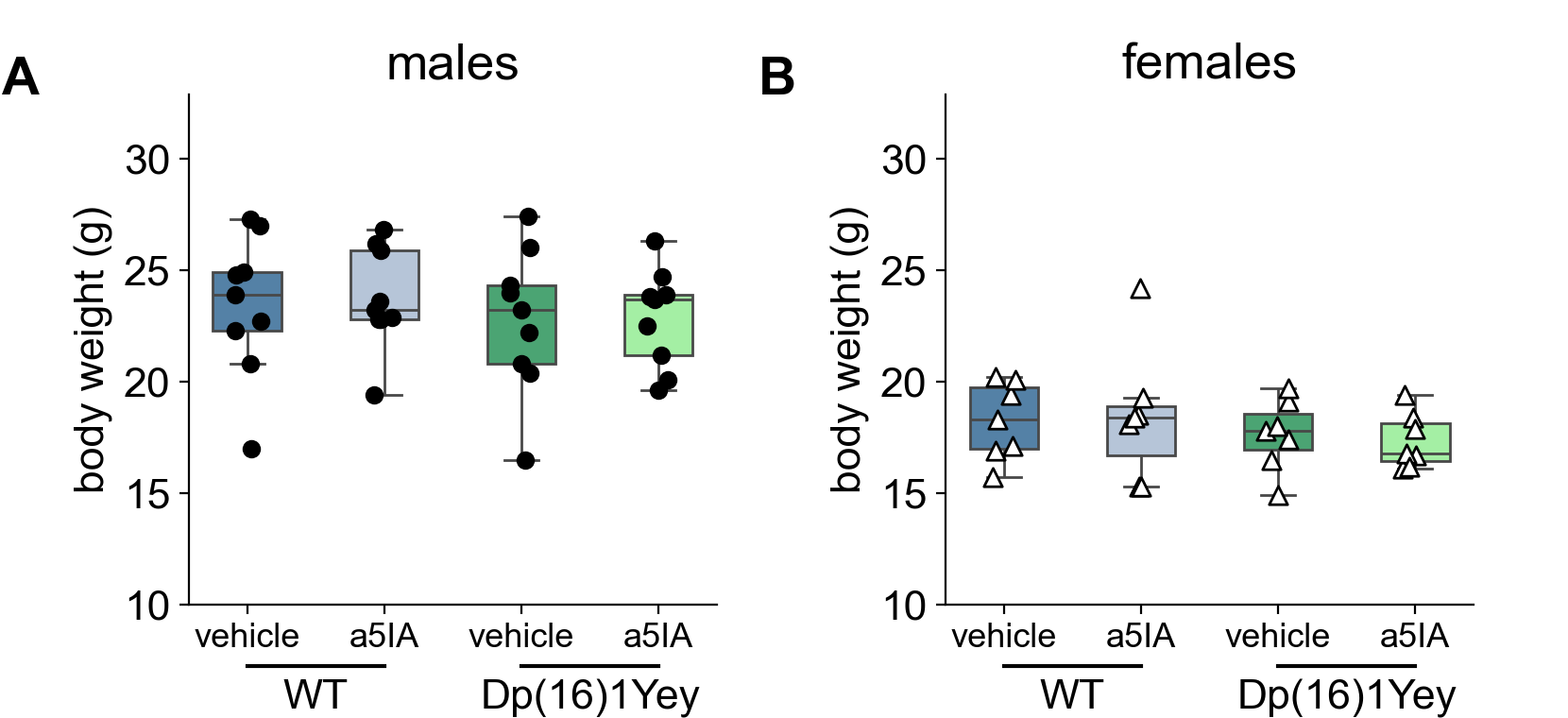

**Supplementary Figure S3: α5IA did not impact body weight of the mice. Body weight of (A) male and (B) female mice at the end of the pattern dissociation task.**

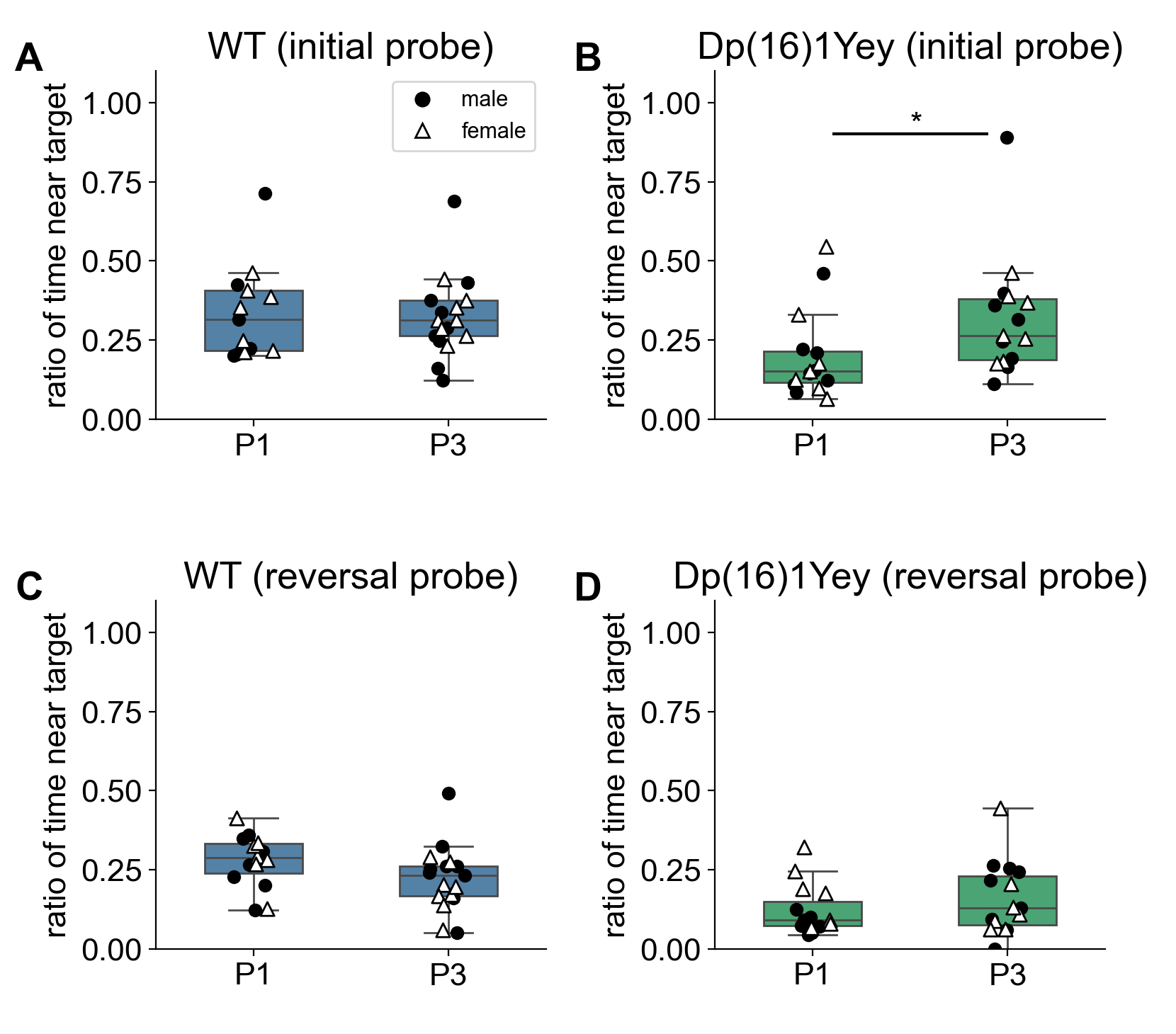

**Supplementary Figure S4**: **Dp(16)1Yey mice might be sensitive to cognitive stimulation**. Ratio of time spent near the escape hole during the probe phase of the Barnes maze for (A) WT vehicle and (B) Dp(16)1Yey vehicle mice. Mice from P1 pipeline were naïve to any cognitive test. Mice from the P3 pipeline had performed the touchscreen beforehand. n (WT P1 = 13, WT P3 = 17, Dp(16)1Yey P1 = 15, Dp(16)1Yey P3 = 15). u-test: *: p<0.05 (WT from P1 vs WT vehicle from P3: U=110, p=1; Dp(16)1Yey from P1 vs Dp(16)1Yey vehicle from P3: U=53, p=0.014). Ratio of time spent near the escape hole during the reversal probe phase of the Barnes maze for (A) WT vehicle and (B) Dp(16)1Yey vehicle mice. n (WT P1 = 13, WT P3 = 17, Dp(16)1Yey P1 = 15, Dp(16)1Yey P3 = 15). (WT from P1 vs WT vehicle from P3: t(28)=1.600 p=0.121; Dp(16)1Yey from P1 vs Dp(16)1Yey vehicle from P3: U= 53, p=0.340).
