## Supplementary Methods for "Differential efficacy of α5IA in the Dp(16)1Yey mouse model of Down syndrome: implications for translational research"

### Identification and genotyping procedure

Genotyping and marking were performed either by taking the tips of the finger at 1 week of age or by collecting ear punches at weaning at 4 weeks of age. The genotyping was done using the following primers: Dp16 Yey 1R (GTC AGT GGT TGT GAC TTG TG), Dp16 Yey 1F (TAT TAG GAC AAG GCT GGT GG), Dp16 Yey CI For (CTA GGC CAC AGA ATT GAA AGA TCT), Dp16 Yey CI Rev (GTA GGT GGA AAT TCT AGC ATC ATC C). The program of amplification was: an initial denaturation at 95°C for 2 min, then 35 cycles of amplification (denaturation at 95°C for 30 sec, amplification at 60°C for 30 sec, elongation at 72°C for 30 sec), and final elongation at 72°C for 7 min.

### Procedures for behavioral tests

#### Barnes maze

The Barnes maze test was performed on a home-made white PVC circular board following the blueprint from conductscience.com (with a diameter of 92 cm and 20 holes at the periphery, each with a diameter of 5 cm). The platform was 95 cm above the ground. The maze was illuminated with 130 lux at the center, and visual cues were placed on the walls. One hole was connected to an escape box. The location of the escape hole (in the room referential) was the same for all mice. Between each trial, all the holes and the escape box were cleaned using water mixed with detergent. The maze was rotated seven holes in a counterclockwise motion to prevent mice using olfactory clues instead of spatial ones to find the escape hole. The escape box was replaced at the correct position. Video capture and primary tracking were done using Ethovision XT software (version 14; Noldus, Wageningen, the Netherlands).

Each trial started similarly. The mouse was placed under an opaque bucket (height 20 cm; diameter 10 cm) in the middle of the arena. The timer was launched when the bucket was removed. On the first day, the mice were introduced in the arena for 3 min of free exploration. An escape hole was also present in this habituation phase, but its position differed from the one used during the learning phase. At the end of the time, if the mouse was still on the table, she was gently guided toward the escape box by blocking progressively all other ways. Then, during the six days of the learning phase (second to seventh days), each mouse performed daily three trials of 3 min with a 30-min intertrial break. The recording ended when the mouse went inside the escape box or at the end of the 3 min (in this case, the mouse was gently guided with the hand into the escape hole). For the pipeline with the treatment (P3), we performed two trials per day instead of three trials per day. On the eighth day, each mouse performed one 1 min 30 s probe trial, while no escape box was connected to the maze. The next day, we started a reversal phase that lasted three days (ninth to eleventh days). The mice performed daily three trials of 3 min with a 30-min intertrial break while the treated cohort performed only two trials per day. This time, the escape box was connected to the diametrically opposite hole compared to the learning phase. Finally, on the twelfth day, we performed again a 1 min 30 s probe test. We assessed the time spent in the correct quarter and next to the target hole during the two probe phases.

#### Rotarod

We performed a rotarod test to evaluate motor functions related to the somatosensory cortex. We used the protocol described by Aziz and colleagues (Aziz et al., 2018). Briefly, mice were evaluated over two days. On the first day, eight trials of 120 s with a 15 min intertrial break were performed by each mouse. First, the mice were habituated to the apparatus during two trials at 16 rotations per minute (RPM), and then they were evaluated at different speeds for two trials each time (16, 24, and 32 RPM). During the second day, the mice performed two trials with progressively increasing speed from 4 to 40 RPM over 5 min. The apparatus was cleaned between trials with 70% ethanol. The latency to fall was recorded if the animal fell off the rod or made three passive turns around the rod during the trial.

#### Y-maze

To evaluate working memory, we used the Y-maze. Mice were placed in a three arms Y-shaped maze (an angle of 120° separated each arm that measured 40*9*16 cm) illuminated at 130 lux, and with different graphical patterns on each arm. They explored the maze freely for 8 min. During this period, we counted the number of arm entries (i.e., the entire body of the mouse reached half of the length of the arm) and the sequence of arm visits to determine the proportion of spontaneous alternations and the number of returns in the arm. A spontaneous alternation was described as follows: the mouse explored the three different arms in a row. Spontaneous alternation ratio is calculated as the number of spontaneous alternations divided by the number of possible alternations (number of arm entries - 2). The maze was cleaned with 70% ethanol between two mice.

#### Pattern dissociation task

To assess learning capacity in Dp(16)1Yey mice, we used the pattern dissociation paradigm on Bussey-Saksida touch screen systems (80614A, Lafayette lifesciences, Loughborough, UK). The goal was to assess learning specific task with a system derived from the CANTAB (CAmbridge Neuropsychological Test Automated Battery) evaluations that are used with patients (Nithianantharajah et al., 2015). We used acidification of the water in the housing cage to avoid food or water privation while maintaining high motivation (Jehl et al., 2025). The test enclosure was cleaned between two sessions with 70% ethanol, followed by detergent mixed with water. The screen was cleaned with humidified tissues soaked in detergent water mix.

First, during one week, mice were habituated to the reward (half-skimmed milk and strawberry syrup; 12% sugar) with free access for one hour per day. To increase the motivational value of the reward, we gradually acidified the drinking water in the housing cages during weekdays, while providing standard drinking water on weekends. In the first week, the water pH was adjusted to 4 for the initial two days, followed by a further reduction to pH 3 for the remainder of the week. To monitor weight loss, we compared their daily body weight to their initial body weight measured on the first day.

During the following week, the mice were trained to use the touch screen. We used three schedules for that. The first one was for learning the location of the reward. For that, a stimulus appeared on the screen for 20 s. If the mice interacted, they received a standard reward (20 µL) with a noise and a light appearing in the feeder. If the mice did not interact with the screen, they only received a small reward (5 µL). This schedule included 30 trials (the counter only activated if the mice reacted) and lasted a maximum of 60 min on one day. The second schedule forced the mice to interact with the screen. The stimulus stayed on the screen this time, and there was no reward if the mouse did not interact. We used this schedule two days in a row, once with 30 trials, then with 60 trials; both were 60 min maximum. Finally, we used a schedule where the mice learned to initiate the trial. For that, the feeder was illuminated, and the mouse needed to poke inside to trigger the stimulus on the screen. We used this schedule for two days, with 60 trials that lasted 60 min. During this week, the water pH was 2.8 on the first day, 2.6 for two days, and 2.5 for the last two days.

Then, during four weeks, the mice performed the pattern dissociation task. In this task, the mice needed to learn which picture in a pair was associated with a reward. We used a concentric line picture as the positive picture and a dotted picture for the negative stimulus. To start a trial, the feeder was illuminated. When the mice poked it, the picture pair appeared randomly to avoid side preference. If the mice responded correctly, a sound was broadcasted, the feeder was illuminated, and a 20 µl reward was distributed followed by a 20 s intertrial interval. If the mice chose the image that was not reinforced, the light turned on in the cage for 5 s. In this case, an inter-trial interval of 20 s prevented interaction with the feeder or the screen. Each mouse had one session per day for twenty days. Each session allowed a maximum of 60 trials or 60 min. The percent of success and the trial number were recorded. During this period, the water pH decreased by 0.1 per week, from 2.4 to 2.1. For the treatment pipeline (P3), we reduced the pattern dissociation phase to two weeks following the timing described previously (Jehl et al., 2025).

We evaluated the percent of success per session during all the learning period, the number of sessions to reach 75 percent of success over the 60 trials of the session (arbitrary threshold where we consider that the mouse had understood the rule), the time to perform the 60 trials during the session, and the percent of success during the 10^th^ session (P2 and P3) and the 20^th^ session (P3). To assess the global success of the mice during the learning period, we calculated the area under the learning curve (AUC). We used this information to summarize the learning capacity of the tested group in Gdaphen (Muñiz Moreno et al., 2023).

#### Novel object recognition

To assess declarative memory, we use the novel object recognition test. During the first day, the mice were habituated to an opaque round arena (diameter: 1 m; height: 40 cm) for 10 min at 130 lux. We recorded the total distance traveled to evaluate activity in a new environment.

During the second day, a pair of similar objects (flask/flask or bottle/bottle) was introduced into the same round arena. We used two types of objects: a T25 plastic cell culture flask (painted in black from inside; 4 x 8 x 2 cm; (353018, Corning SAS, SAMOIS-SUR-SEINE, FR) and a cleaned glass bottle (emptied, autoclaved then filled with sand; height: 11.5 cm; diameter: 4 cm (Safran thread, Saint Lucie 1885, CREIL, FR). We fixed the object in the arena using adhesive putty. Both objects had similar size and global circumference and could be climbed on by the mice. We verified that mice did not display significant spontaneous preference between these two objects during this acquisition phase (Supplementary Figure S1A & B). Mice were left to freely explore the pair of objects for 10 min. We registered the time spent exploring the objects (nose pointing toward the object within 3 cm), excluding the time spent on top of the object. We calculated a side ratio as time_exploring_right_object / (time_exploring_right_object + time_exploring_left_object).

The following day, mice were again placed in the same arena, in which we placed the familiar object (i.e., the one that had been presented the day before) in the left or right side randomly and an unfamiliar object (i.e., the novel one that had not been presented the day before). We recorded the time spent exploring the objects as described above. We calculated the proportion of exploration of the novel object as time_exploring_novel_object / (time_exploring_novel_object + time_exploring_familiar_object). In all the phases, the arena and the objects were cleaned using 70% ethanol between two mice.
