## Supplementary tables for "Differential efficacy of α5IA in the Dp(16)1Yey mouse model of Down syndrome: implications for translational research"

**Supplementary Table S1:** Computation of the statistical power and estimation of sample size for each test of the behavioral characterization.

| **Test** | **Mean WT** | **SD WT** | **Mean Dup** | **SD Dup** | **Effect size** | **Individuals needed per group** |
| --- | --- | --- | --- | --- | --- | --- |
| **Y-maze wt vs dup** | 0,64 | 0,08 | 0,59 | 0,09 | 0,59 | 47 |
| **Ymaze vs 50% Wt** | 0,64 | 0,08 | / | / | 1,75 | 6 |
| **Ymaze vs 50% Dup** | / | / | 0,59 | 0,09 | 1 | 13 |
| **Barnes maze (reverse hole)** | 0,28 | 0,08 | 0,12 | 0,08 | 2 | 6 |
| **Touchscreen** | 1522,9 | 127,58 | 1185,2 | 185,85 | 2,12 | 5 |
| **Rotarod** | 248,0 | 48,3 | 174,4 | 57,2 | 1,39 | 10 |

**Supplementary Table S2**: Results of the two ways ANOVA (genotype, sex) of the characterization cohorts (SS: sum of squares, df: degree of freedom, Geno: genotype)

| **Y-maze / spontaneous alternation:** | | | | | | |
| --- | --- | --- | --- | --- | --- | --- |
|  | **Source of variability** | **SS** | **df** | **F** | **p** | **Significance** |
|  | **Geno** | 0,041302 | 1 | 5,791408 | 0,019311 | * |
|  | **Sex** | 0,000693 | 1 | 0,097226 | 0,756302 |  |
|  | **Geno : Sex** | 0,002381 | 1 | 0,333926 | 0,565593 |  |
|  | **Residual error** | 0,413636 | 58 | / | / |  |
|  | **Total** | 0,458012 | 61 | / | / |  |
| **Y-maze / arm entries:** | | | | | | |
|  | **Source of variability** | **SS** | **df** | **F** | **p** | **Significance** |
|  | **Geno** | 0,340858 | 1 | 0,003957 | 0,950061 |  |
|  | **Sex** | 234,717093 | 1 | 2,72451 | 0,104224 |  |
|  | **Geno : Sex** | 160,965882 | 1 | 1,868433 | 0,17693 |  |
|  | **Residual error** | 4996,71201 | 58 | / | / |  |
|  | **Total** | 5392,73584 | 61 | / | / |  |
| **Barnes maze / time spent near the escape (initial probe):** | | | | | | |
|  | **Source of variability** | **SS** | **df** | **F** | **p** | **Significance** |
|  | **Geno** | 0,127343 | 1 | 5,760284 | 0,024507 | * |
|  | **Sex** | 0,000119 | 1 | 0,005377 | 0,942152 |  |
|  | **Geno : Sex** | 0,003669 | 1 | 0,16596 | 0,687338 |  |
|  | **Residual error** | 0,53057 | 24 | / | / |  |
|  | **Total** | 0,661701 | 27 | / | / |  |
| **Barnes maze / time spent in the correct quarter (initial probe):** | | | | | | |
|  | **Source of variability** | **SS** | **df** | **F** | **p** | **Significance** |
|  | **Geno** | 0,023448 | 1 | 1,343462 | 0,257827 |  |
|  | **Sex** | 0,004154 | 1 | 0,237983 | 0,630093 |  |
|  | **Geno : Sex** | 0,018966 | 1 | 1,086647 | 0,307603 |  |
|  | **Residual error** | 0,418885 | 24 | / | / |  |
|  | **Total** | 0,465453 | 27 | / | / |  |
| **Barnes maze / time spent near the escape (reversal probe):** | | | | | | |
|  | **Source of variability** | **SS** | **df** | **F** | **p** | **Significance** |
|  | **Geno** | 0,173597 | 1 | 29,375341 | 0,000013 | *** |
|  | **Sex** | 0,026328 | 1 | 4,455139 | 0,04497 | * |
|  | **Geno : Sex** | 0,006049 | 1 | 1,023666 | 0,321344 |  |
|  | **Residual error** | 0,14774 | 25 | / | / |  |
|  | **Total** | 0,353714 | 28 | / | / |  |
| **Barnes maze / time spent in the correct quarter (reversal probe):** | | | | | | |
|  | **Source of variability** | **SS** | **df** | **F** | **p** | **Significance** |
|  | **Geno** | 0,079138 | 1 | 9,070948 | 0,005869 | ** |
|  | **Sex** | 0,008466 | 1 | 0,970376 | 0,334025 |  |
|  | **Geno : Sex** | 0,030032 | 1 | 3,442289 | 0,075376 |  |
|  | **Residual error** | 0,218108 | 25 | / | / |  |
|  | **Total** | 0,335744 | 28 | / | / |  |
| **Pattern dissociation / Average success 10^th^ session:** | | | | | | |
|  | **Source of variability** | **SS** | **df** | **F** | **p** | **Significance** |
|  | **Geno** | 6426.496718 | 1 | 32.657987 | 4.197644E-07 | *** |
|  | **Sex** | 82.797831 | 1 | 0.420760 | 5.191631E-01 |  |
|  | **Geno : Sex** | 129.023995 | 1 | 0.655670 | 4.214602e-01 |  |
|  | **Residual error** | 11216.561143 | 57 | / | / |  |
|  | **Total** | 17854,8797 | 60 | / | / |  |
| **Pattern dissociation / Average success 20^th^ session:** | | | | | | |
|  | **Source of variability** | **SS** | **df** | **F** | **p** | **Significance** |
|  | **Geno** | 3775.865042 | 1 | 25.886905 | 0.000004 | *** |
|  | **Sex** | 23.691763 | 1 | 0.162428 | 0.688413 |  |
|  | **Geno : Sex** | 1.598908 | 1 | 0.010962 | 0.916976 |  |
|  | **Residual error** | 8459.882282 | 57 | / | / |  |
|  | **Total** | 12261,038 | 60 | / | / |  |

| **Novel object recognition / exploration index (acquisition):** | | | | | | |
| --- | --- | --- | --- | --- | --- | --- |
|  | **Source of variability** | **SS** | **df** | **F** | **p** | **Significance** |
|  | **Geno** | 0,011746 | 1 | 1,477858 | 0,235033 |  |
|  | **Sex** | 0,008603 | 1 | 1,08247 | 0,307724 |  |
|  | **Geno : Sex** | 0,003459 | 1 | 0,435191 | 0,515254 |  |
|  | **Residual error** | 0,206646 | 26 | / | / |  |
|  | **Total** | 0,230454 | 29 | / | / |  |
| **Novel object recognition / Recognition index (test):** | | | | | | |
|  | **Source of variability** | **SS** | **df** | **F** | **p** | **Significance** |
|  | **Geno** | 0,0012 | 1 | 0,118859 | 0,733049 |  |
|  | **Sex** | 0,00024 | 1 | 0,023735 | 0,878751 |  |
|  | **Geno : Sex** | 0,011551 | 1 | 1,143756 | 0,294689 |  |
|  | **Residual error** | 0,262585 | 26 | / | / |  |
|  | **Total** | 0,275576 | 29 | / | / |  |
| **Rotarod / acceleration:** | | | | | | |
|  | **Source of variability** | **SS** | **df** | **F** | **p** | **Significance** |
|  | **Geno** | 42703,6519 | 1 | 13,958214 | 0,000849 | *** |
|  | **Sex** | 3110,63281 | 1 | 1,016749 | 0,321926 |  |
|  | **Geno : Sex** | 2780,28584 | 1 | 0,908771 | 0,348599 |  |
|  | **Residual error** | 85662,9841 | 28 | / | / |  |
|  | **Total** | 134257,555 | 31 | / | / |  |

**Supplementary Table S3**: Results of the three ways ANOVA (genotype, treatment, sex) of the treatment cohorts (SS: sum of squares, df: degree of freedom, Geno: genotype, Treat: treatment)

| **Y-maze / spontaneous alternation:** | | | | | | |
| --- | --- | --- | --- | --- | --- | --- |
|  | **Source of variability** | **SS** | **df** | **F** | **p** | **Significance** |
|  | **Geno** | 0.030275 | 1 | 3.775980 | 0.057026 |  |
|  | **Treat** | 0.001992 | 1 | 0.248433 | 0.620133 |  |
|  | **Sex** | 0.000812 | 1 | 0.101277 | 0.751486 |  |
|  | **Geno: Treat** | 0.000590 | 1 | 0.073638 | 0.787110 |  |
|  | **Geno: Sex** | 0.018465 | 1 | 2.302981 | 0.134751 |  |
|  | **Treat: Sex** | 0.014625 | 1 | 1.824030 | 0.182267 |  |
|  | **Geno: Treat: Sex** | 0.020272 | 1 | 2.528410 | 0.117443 |  |
|  | **Residual error** | 0.448997 | 56 | / | / |  |
|  | **Total** | 0,536028 | 63 | / | / |  |
| **Y-maze / arm entries:** | | | | | | |
|  | **Source of variability** | **SS** | **df** | **F** | **p** | **Significance** |
|  | **Geno** | 76,5625 | 1 | 1,310971 | 0,257089 |  |
|  | **Treat** | 506,25 | 1 | 8,668462 | 0,004707 | ** |
|  | **Sex** | 305,580357 | 1 | 5,232418 | 0,025969 | * |
|  | **Geno : Treat** | 12,25 | 1 | 0,209755 | 0,648731 |  |
|  | **Geno : Sex** | 196,453373 | 1 | 3,363849 | 0,071957 |  |
|  | **Treat : Sex** | 42,099206 | 1 | 0,72086 | 0,399477 |  |
|  | **Geno: Treat: Sex** | 118,765873 | 1 | 2,033615 | 0,159405 |  |
|  | **Residual error** | 3270,47619 | 56 | / | / |  |
|  | **Total** | 4528,4375 | 63 | / | / |  |
| **Barnes maze / time spent near the escape hole (probe):** | | | | | | |
|  | **Source of variability** | **SS** | **df** | **F** | **p** | **Significance** |
|  | **Geno** | 0,008747 | 1 | 0,334909 | 0,565103 |  |
|  | **Treat** | 0,039751 | 1 | 1,521915 | 0,222483 |  |
|  | **Sex** | 0,012382 | 1 | 0,474064 | 0,493966 |  |
|  | **Geno: Treat** | 0,036205 | 1 | 1,386185 | 0,24403 |  |
|  | **Geno: Sex** | 0,000507 | 1 | 0,019428 | 0,889648 |  |
|  | **Treat: Sex** | 0,003678 | 1 | 0,140815 | 0,70889 |  |
|  | **Geno: Treat: Sex** | 0,000159 | 1 | 0,00608 | 0,938125 |  |
|  | **Residual error** | 1,462651 | 56 | / | / |  |
|  | **Total** | 1,56408 | 63 | / | / |  |

| **Barnes maze / time spent in the correct quarter (initial probe):** | | | | | | |
| --- | --- | --- | --- | --- | --- | --- |
|  | **Source of variability** | **SS** | **df** | **F** | **p** | **Significance** |
|  | **Geno** | 0,060777 | 1 | 2,434826 | 0,124301 |  |
|  | **Treat** | 0,029899 | 1 | 1,197801 | 0,278447 |  |
|  | **Sex** | 0,015962 | 1 | 0,63946 | 0,427285 |  |
|  | **Geno : Treat** | 0,045411 | 1 | 1,819245 | 0,182832 |  |
|  | **Geno : Sex** | 0,003029 | 1 | 0,121349 | 0,728883 |  |
|  | **Treat : Sex** | 0,000828 | 1 | 0,033153 | 0,856177 |  |
|  | **Geno: Treat: Sex** | 0,000033 | 1 | 0,001328 | 0,971064 |  |
|  | **Residual error** | 1,397838 | 56 | / | / |  |
|  | **Total** | 1,553777 | 63 | / | / |  |
| **Barnes maze / time spent near the escape hole (reversal probe):** | | | | | | |
|  | **Source of variability** | **SS** | **df** | **F** | **p** | **Significance** |
|  | **Geno** | 0,056494 | 1 | 7,257684 | 0,009299 | ** |
|  | **Treat** | 0,000005 | 1 | 0,000649 | 0,979769 |  |
|  | **Sex** | 0,002672 | 1 | 0,343261 | 0,560307 |  |
|  | **Geno: Treat** | 0,006642 | 1 | 0,853231 | 0,359603 |  |
|  | **Geno: Sex** | 0,031697 | 1 | 4,072006 | 0,0484 | * |
|  | **Treat: Sex** | 0,00347 | 1 | 0,445763 | 0,507096 |  |
|  | **Geno: Treat: Sex** | 0,000784 | 1 | 0,100753 | 0,752108 |  |
|  | **Residual error** | 0,435906 | 56 | / | / |  |
|  | **Total** | 0,53767 | 63 | / | / |  |
| **Barnes maze / time spent in the correct quarter (reversal probe):** | | | | | | |
|  | **Source of variability** | **SS** | **df** | **F** | **p** | **Significance** |
|  | **Geno** | 0,010289 | 1 | 0,460625 | 0,500127 |  |
|  | **Treat** | 0,004959 | 1 | 0,221997 | 0,639353 |  |
|  | **Sex** | 0,019999 | 1 | 0,895303 | 0,348109 |  |
|  | **Geno : Treat** | 0,002661 | 1 | 0,119138 | 0,731265 |  |
|  | **Geno : Sex** | 0,017714 | 1 | 0,793017 | 0,376999 |  |
|  | **Treat : Sex** | 0,000063 | 1 | 0,00282 | 0,957839 |  |
|  | **Geno: Treat: Sex** | 0,001268 | 1 | 0,056769 | 0,812548 |  |
|  | **Residual error** | 1,250895 | 56 | / | / |  |
|  | **Total** | 1,307848 | 63 | / | / |  |

| **Pattern dissociation / Average success 10th session:** | | | | | | |
| --- | --- | --- | --- | --- | --- | --- |
|  | **Source of variability** | **SS** | **df** | **F** | **p** | **Significance** |
|  | **Geno** | 5018.242513 | 1 | 28.838697 | 0.000002 | *** |
|  | **Treat** | 1.916762 | 1 | 0.011015 | 0.916788 |  |
|  | **Sex** | 57.871715 | 1 | 0.332576 | 0.566457 |  |
|  | **Geno: Treat** | 2.808060 | 1 | 0.016137 | 0.899370 |  |
|  | **Geno: Sex** | 367.558081 | 1 | 2.112273 | 0.151703 |  |
|  | **Treat: Sex** | 74.876053 | 1 | 0.430296 | 0.514530 |  |
|  | **Geno: Treat: Sex** | 7.967058 | 1 | 0.045785 | 0.831345 |  |
|  | **Residual error** | 9744.600478 | 56 | / | / |  |
|  | **Total** | 15275.8407 | 63 |  |  |  |
| **Rotarod / acceleration:** | | | | | | |
|  | **Source of variability** | **SS** | **df** | **F** | **p** | **Significance** |
|  | **Geno** | 90450,5625 | 1 | 52,538004 | 1,35E-09 | *** |
|  | **Treat** | 8602,5625 | 1 | 4,996779 | 2,94E-02 | * |
|  | **Sex** | 11,464534 | 1 | 0,006659 | 9,35E-01 |  |
|  | **Geno: Treat** | 682,515625 | 1 | 0,396438 | 5,31E-01 |  |
|  | **Geno: Sex** | 1,834325 | 1 | 0,001065 | 9,74E-01 |  |
|  | **Treat: Sex** | 2003,21528 | 1 | 1,163563 | 2,85E-01 |  |
|  | **Geno: Treat: Sex** | 1382,52009 | 1 | 0,803034 | 3,74E-01 |  |
|  | **Residual error** | 96410,8095 | 56 | / | / |  |
|  | **Total** | 199545,484 | 63 | / | / |  |

**Supplementary Table S4:** Name and definition of the variables used to conduct the GDAPHEN multivariate analysis.

| **abbreviation** | **full name** | **function evaluated** |
| --- | --- | --- |
| weight | Body weight measurement at the end of the pattern dissociation test | animal general state |
| returns_in_arms | Number of times a mouse did not choose a new arm during the Y-maze test | anxiety |
| AUC_touchscreen | Area under the curve of the success rate during the learning of the pattern dissociation test | learning |
| threshold_touchscreen | Number of sessions needed to reach the threshold | learning |
| acceleration_rotarod | Latency to fall during the rotarod's acceleration paradigm | motor function |
| number_arms | Number of arms explored during the Y-maze test | motor function |
| rpm_16 | Latency to fall during the rotarod test at 16 rotations per minute | motor function |
| rpm_24 | Latency to fall during the rotarod test at 24 rotations per minute | motor function |
| rpm_32 | Latency to fall during the rotarod test at 32 rotations per minute | motor function |
| time_hole_prob | Ratio of time spent near the correct hole during the initial probe phase of the Barnes maze | spatial memory |
| time_quarter_prob | Ratio of time spent in the correct quarter during the initial probe phase of the Barnes maze | spatial memory |
| time_hole_rev | Ratio of time spent near the correct hole during the reversal probe phase of the Barnes maze | spatial memory / cognitive flexibility |
| time_quarter_rev | Ratio of time spent in the correct quarter during the reversal probe phase of the Barnes maze | spatial memory / cognitive flexibility |
| spontaneous_alternation | Percent of spontaneous alternations during the Y-maze | working memory |
